# Micro-CT Analysis of a Needle to Nerve Approach for Interfascicular Peripheral Nerve Stimulation in Porcine Nerves

**DOI:** 10.1101/2025.11.19.687256

**Authors:** Margaux A. Randolph, Vlad I. Marcu, Constantinos Tsiptsis, Andrew J. Shoffstall, Jonathan Z. Baskin, Dustin J. Tyler

## Abstract

**Motivation:** Peripheral nerve interfaces typically have a tradeoff between selectivity and invasiveness. Interfascicular interfaces placed within the peripheral nerve, but outside of the fascicles preserving the perineurium are a promising avenue in balancing this tradeoff. This study quantifies electrode placement using novel tools designed to deliver flexible microwires interfascicularly in a porcine model.

**Methods:** Seven Yorkshire pigs were implanted with Minimally invasive interfascicular Nerve Stimulation (MiiNS) arrays in the nerves of the brachial plexus resulting in a total of ten implanted nerve samples. High-resolution micro-computed tomography (micro-CT) of implanted nerves stained with phosphotungistic acid (PTA) showed contact placement relative to fascicular anatomy and all other contacts. The analysis also examined the relationship between microwire trajectory angle and fascicle traversal. Further, in a subset of samples (n=4), hematoxylin and eosin (H&E) histological analysis was performed after micro-CT imaging. These histological images were coregistered with the micro-CT to validate the positional information of the micro-CT.

**Results:** Across the 56 total MiiNS microwire placements, 84% were interfascicular, while 16% were intrafascicular, characterized by traversal through a fascicle. Contacts were broadly distributed throughout the nerve’s cross section and the majority (79%) were in the central half of the nerve’s cross section (R>0.707), with no evidence of angular clustering in any single direction. On average, nearest neighbor distances between contacts measured 2256.26±1760.28 µm in 3D and 730.71 ±564.83 µm transversely, with implants spanning 10.3 ± 4.3 mm along length of nerve. The angle of wire trajectory into the nerve was correlated with fascicle traversal, with steeper angles of insertion associated with more instances of fascicle traversal. Histological analysis corroborated the fascicular borders and wire placements found on micro-CT and demonstrated perineurium integrity.

**Significance:** Interfascicular implantation can reliably access both central and distributed regions of the nerve, while not being confined to a single plane, enabling access to regions which have been historically challenging to stimulate effectively. The observed nearest-neighbor transverse spacing between contacts is within the range of reported ideal values for selective activation. The MiiNS placement characteristics show its potential as an effective peripheral nerve interface alternative which achieves distributed contact placement, including in the center of the nerve volume, while remaining predominantly outside of the perineurium.

## Introduction

Peripheral nerve stimulation (PNS) has been implemented for various clinical applications including alleviation of chronic neuropathic pain, sensory restoration following amputation and a wide range of autonomic therapies[1–3]. Peripheral nerve stimulation is an expanding field of study, with new nerve targets and treatment approaches emerging regularly. The therapeutic benefit of peripheral nerve interfaces across applications often relies on the selectivity of the interface, which refers to its capability to activate specific axonal populations while avoiding the activation of others. Selectivity is especially important in neuroprosthetic applications, such as the restoration of motor or sensory functions. However, interfaces that achieve a higher selectivity are generally more invasive[4,5]. Invasiveness refers to the extent of surgical intervention required for implantation and long-term structural or functional disruption of nerve tissue resulting from chronic implantation. This tradeoff between selectivity and invasiveness remains an active area of development for the implementation of clinically optimal neural interfaces.

### Current state of Peripheral nerve interfaces

Historically, peripheral nerve interfaces across neuroprosthetic applications have developed towards the selective activation of small populations of axons. To achieve this, interfaces are designed to be placed at various positions throughout the nerve. Circumneural interfaces are placed around the nerve and demonstrate selective activation, especially when the interface causes gentle reshaping of the nerve, as demonstrated in the composite flat interface nerve electrode (CFINE)[6–8]. These interfaces do not penetrate the perineurium or the epineurium but require circumneural dissection for implantation. Currently, circumneural electrodes are used following amputation for sensory and motor restoration, and for the management of postamputation pain[8–10]. However, in large, poly-fasciculated nerves of human subjects, circumneural electrodes cannot selectively activate central axonal populations, even with reshaping, given the practical geometrical limits of reshaping and the desire to maintain the applied pressure on the nerve below threshold safety values [11]. Alternatively, intrafascicular electrodes aim to increase selectivity by interfacing with the axonal population directly, within the nerve and fascicles[12–14]. Intrafascicular interfaces restore sensory and motor function and selectively activate the nerves[13,15,16]. However, in rupturing the blood nerve barrier, morphological changes such as axon loss, and changes to axon diameter distribution have been observed across several variations of intrafascicular interfaces[17–22].

### Minimally invasive interfascicular nerve stimulation (MiiNS)

may address the key limitations of existing peripheral nerve interfaces. Both circumneural and intrafascicular electrodes have a limited ability to access central and distributed regions within poly-fasciculated nerves. This limitation may be addressed by placing contacts inside the nerve, between the fascicles, which also preserves the integrity of the blood-nerve barrier. The concept of an interfascicular peripheral nerve electrode was first introduced with the slowly penetrating interfascicular nerve electrode (SPINE)[23]. Kopakka et al first studied the evoked stimulation selectivity of directional interfascicular electrodes wherein contacts were placed acutely in the sciatic nerve of a rabbit[11]. This approach demonstrated selective stimulation of downstream muscles, motivating further study of interfascicular electrode configurations. However, the rabbit sciatic only had 2-3 fascicles at the reported site of implantation, making extrapolation of results to human nerves uncertain. The spatial distribution of electrode contacts within the nerve was not quantified in this study and the extent of fascicular penetration along wire trajectories was not reported.

### Mitigating Insertion Challenges of Peripheral Nerve Interfaces

Creating a neural interface that is as small and flexible as feasibly possible is important in mitigating the inflammatory response[24]. Insertion of flexible materials into neural tissue remains a significant challenge because of the tendency of these wires to buckle under force before penetration into neural tissue. Various strategies have been explored to address this issue[25]. These include mechanical bracing, transient stiffening materials, ultrasonic-assisted insertion, and the use of mechanically dynamic materials that adapt their stiffness in response to external stimuli[25,26]. Despite these advances, many of these techniques add complexity to the insertion process and have been less explored in the periphery. Flexible intrafascicular interfaces such as carbon nano tubes, LIFE and TIME electrodes, rely on tying or winding the interface around a sharp needle and passing it through the nerve, then is anchoring via glue[12,27,28]. Often a microscope or magnification is required to effectively thread the device into a fascicle. For rigid intrafascicular interfaces such as the Utah slant array, a pneumatic device is used to penetrate the nerve quickly[29,30]. The insertion process of existing penetrating peripheral nerve electrodes remains a point of friction in the path to translation[31]. Furthermore, recent studies have shown that high-density intrafascicular arrays are also somewhat limited to superficial fascicles[19], or constrained in a single transverse plane in the nerve[15]. Here we explore an additional way to rapidly and reliably place flexible microwire contacts throughout the nerve via a needle-to-nerve approach, which results in a hooked microwire contact in the nerve.

This study quantifies the placement of Minimally Invasive Nerve Stimulation (MiiNS) microwire contacts within poly-fasciculated, clinically relevant porcine nerves, demonstrating interfascicular access and distributed positioning using micro-computed tomography.

## Materials and Methods

### MiiNS Microwire Arrays

Each interfascicular implant contains 3-8 microwire contacts (Fort Wanye Metals, 316 LVM, 50µm diameter, insulated to an outer diameter of 70µm). Each microwire had a contact area of exposed wire which was inserted into a blunt microcannula, resulting a “hooked” configuration for delivery into the nerve (Fig. 1). Care was taken to insert only the distal 2mm of the wire into the microcannula prior to implantation. The microwires were connected to an insulated stranded stainless steel lead wire (Fort Wayne Metals) via conductive epoxy and a mechanical crimp. Lead wires were contained in a silicone tube for routing through the body. Figure 1 demonstrates the concept of the MiiNS interface.

**Figure 1.**
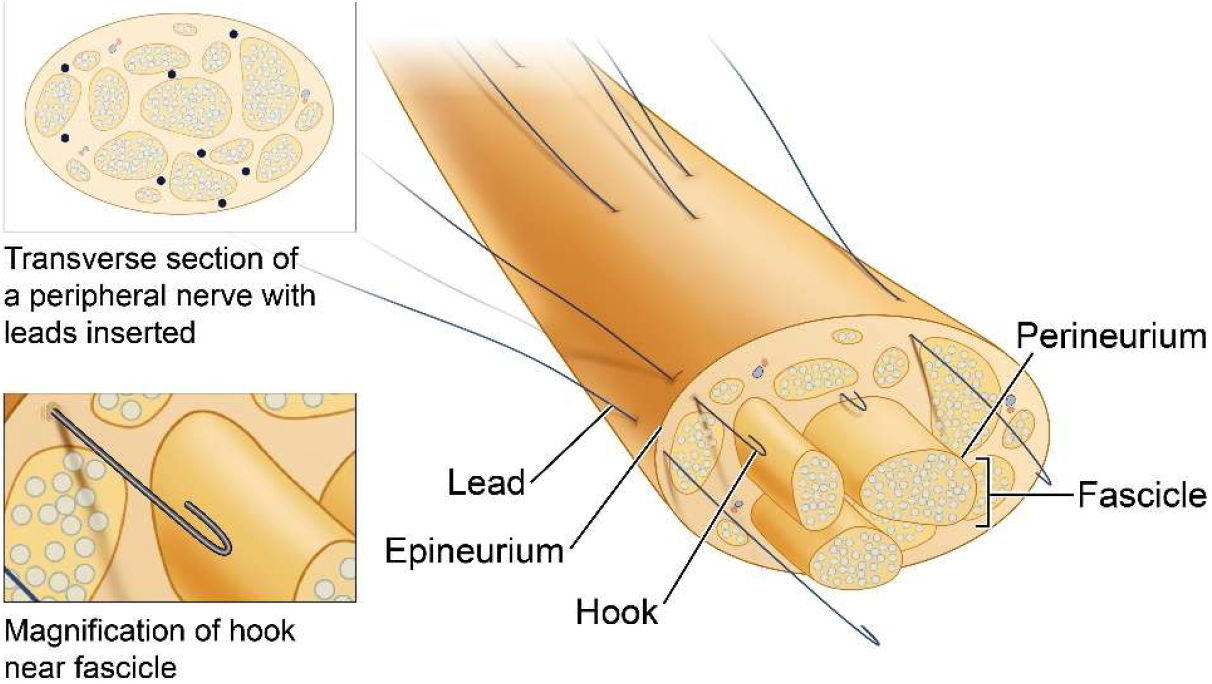
Schematic of MiiNS electrode concept, in a poly-fasciculated peripheral nerve. Multiple hooked microwires are placed between the fascicles within the nerve, penetrating the epineurium, but leaving the perineurium intact (bottom pop out). This configuration allows for access to deep and distributed axonal populations. In the transverse view (top pop-out), contacts are shown to be placed at varying depths and positions between the fascicles, their positions are represented as circles.

#### Animal Cohort

Animal cohort consisted of seven female Yorkshire pigs, with approximate weights ranging 45-95kgs. N=10 total nerves of the brachial plexus (n=7 median and n=3 ulnar, samples 1-10) were implanted with MiiNS and considered for contact placement (Table 1). Nerve samples were obtained immediately following euthanasia.

**Table 1:** Summary of nerve samples, implantation techniques and their treatments.

| Sample | Nerve Type | Implantation Method | Total implanted contacts | Additional Treatment |
| --- | --- | --- | --- | --- |
| 1 | Median | By Hand | 3 |  |
| 2 | Median | By Hand | 7 | Micro-CT Rescan |
| 3 | Median | By Hand | 7 | Micro-CT Rescan |
| 4 | Median | By Hand | 6 | Micro-CT Rescan |
| 5 | Median | Novel Tools | 5 | Histology |
| 6 | Ulnar | Novel Tools | 5 | Histology |
| 7 | Ulnar | Novel Tools | 5 | Histology |
| 8 | Median | Novel Tools | 5 | Histology |
| 9 | Median | Novel Tools | 8 |  |
| 10 | Ulnar | Novel Tools | 5 |  |

#### Surgical Implantation

All surgical instruments were sterilized with either ethylene oxide (EtOH) or autoclave sterilization on a standard instrument cycle. Animals were initially sedated with tiletamine and zolazepam (2-6 mg/kg, IM), before placement of an endotracheal tube, to deliver 2-3% isoflurane to effect throughout the procedure. The pig was placed in a supine position, with the surgical table elevated on the contralateral side to provide increased access to the brachial plexus. Prior to incision, local lidocaine was administered at the incision site to reduce the isoflurane burden. An incision was made on the medial aspect of the upper forelimb 2-5 cm above the medial epicondyle to access the proximal median and ulnar nerves, distal to their respective divergence from the cords. The pectoralis superficialis muscle was divided to the extent necessary to expose the brachial plexus, allowing separation and identification of the vessels and nerves. Bipolar cautery throughout the procedure reduces bleeding while avoiding the application of direct current to nerves of interest. The bicep muscle is useful as an anatomical landmark to identify the median nerve, which is located near the caudal surface of the muscle body and is closely related to the brachial artery. The ulnar nerve was subsequently identified in the caudal direction relative to the median nerve, diverging from the medial cord[32]

#### Surgical Tools for Interfascicular Microwire Implantation

A reliable surgical approach and tools were developed to enable MiiNS implantation. Previous literature on needle geometry recommends 18º tip angles for the successful penetration of the perineurium[33]. The perineurium has a significantly higher penetration force than the epineurium, and this difference is maximized using a blunt probe[34]. A blunt micro-cannula (34g, Cellink) was chosen for implantation of MiiNS to aid in penetration of the epineurium while keeping the perineurium intact.

In samples 1-4, the microwire-loaded blunt microcannulas were affixed to an acrylic shuttle with cyanoacrylate glue and inserted into the nerve via forceps. Once the microcannula was within the nerve, the acrylic shuttle was retracted and removed by hand to leave behind the hooked microwire. In samples 5-10, a novel tool set (Fig. 2) enabled implantation. In these trials, following circumneural exposure, a Neuroretractor (Fig. 2C) was placed beneath the nerve to provide stabilization. The device features flexible bristles that grip the deep surface of epineurium, preventing deflection during oppositional needle insertion on the superficial surface. Then, a spring-loaded inserter delivered a hooked microwire to the nerve via a blunt micro-cannula (Fig. 2A). Upon insertion of the micro-cannula into the nerve, transient, non-damaging pressure was applied to the surface of the nerve using a soft pressure applicator (Fig. 2B). This tool has a gel tip, mechanically locked into a rigid handle, and it temporarily increases the friction between the microwire and the surrounding nerve tissue, to ensure the microwire is reliably left in the nerve. The microcannula was then rapidly retracted via a spring, leaving behind a hooked microwire in the nerve. The rapid retraction of the microcannula enables the microwire to be left behind in the nerve because of the transition from static to dynamic friction at the interface between the microcannula and the microwire. As the cannula moves out of the nerve, the previously tucked distal end of the microwire is released from the cannula, leaving the wire behind in a hooked shape within the nerve. All three devices were 3D printed (Formlabs 3B), with rigid components printed using biomedical resin, and flexible components printed with Elastic 50A (Formlabs).

**Figure 2.**
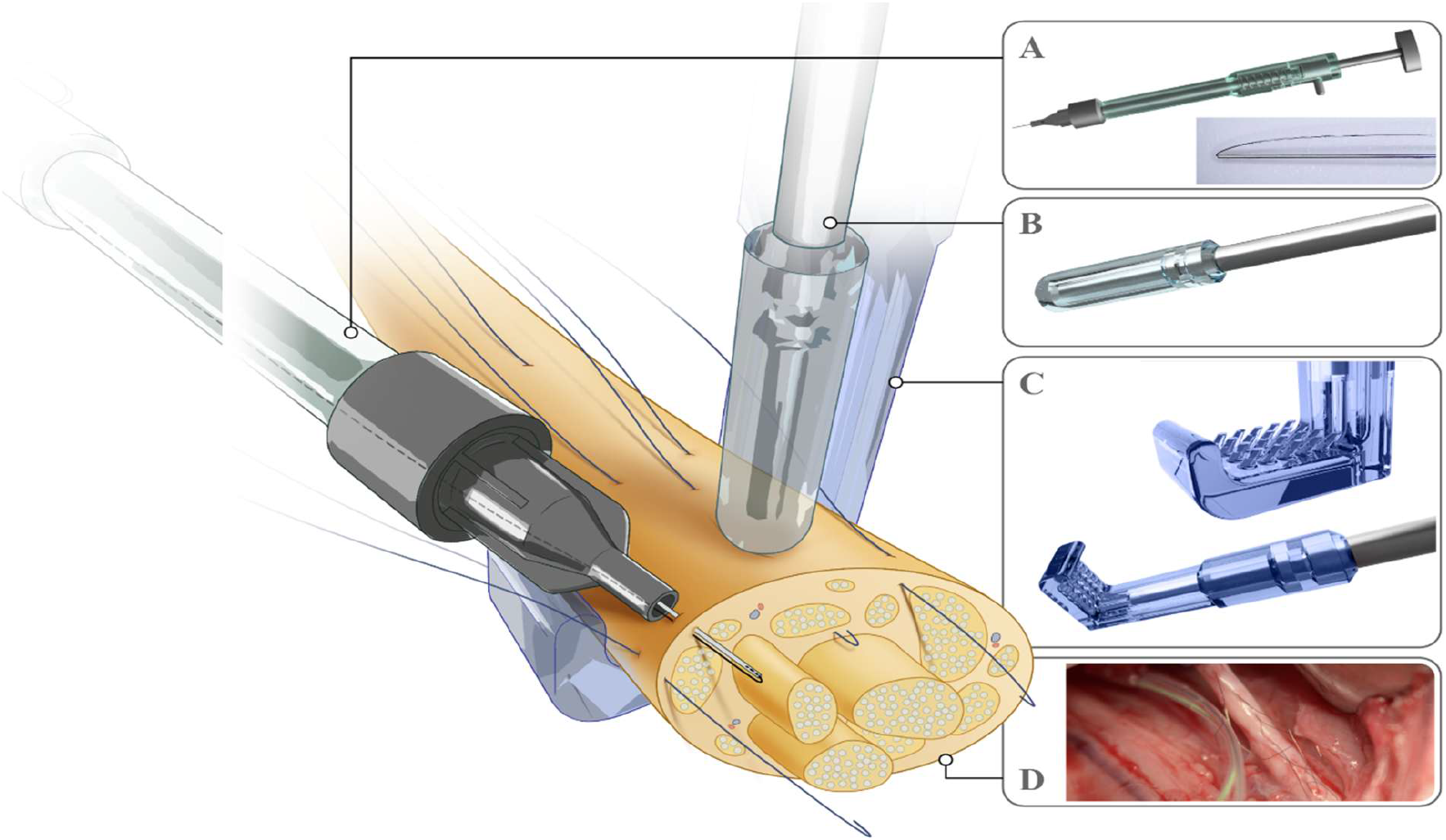
Diagram of novel implantation tools: The spring-loaded inserter (A) with a blunt microcannula attached and pre-loaded with the microwire’s distal end (lower right). The gel tipped pressure applicator (B). The Neuroretractor (C) with flexible bristles for gripping the epineurium following surgical exposure of the nerve. The resulting MiiNS interface with 7 implanted microwire contacts is shown in the porcine median nerve (D).

While the insertion angle was not controlled intraoperatively, the surgeon intuitively placed the microcannula at an oblique angle along the longitudinal axis of the nerve rather than perpendicular to it. The effect of the insertion angle on electrode placement was analysed post hoc. Following successful implant of an electrode, the surgeon discarded the microcannula and attached the inserter to the next microwire-loaded microcannula. Electrode implantation time and success were recorded in two phases: an initial set of four nerve implant trials using manual hand insertion (n = 24 total electrodes) and a subsequent set of five nerve implants using the novel tool set (n=25 total electrodes).

#### Tissue Handling

At the termination of the acute procedure, the implanted nerves were collected and placed directly in 10% neutral buffered formalin (Sigma Aldrich) with a volume at least 50x the total tissue volume for at least 2 days and up to 4 weeks before staining and scanning. Care was taken to minimize the time between the euthanasia and tissue collection to prevent drying and distortion of the neural tissue.

#### Micro-Computed Tomography Staining

To visualize fascicular anatomy within the nerve, phosphotungstic acid (PTA) (3% v/v) (Sigma Aldrich) was prepared with deionized water to increase soft-tissue contrast. Each sample was submerged in 50 mL of PTA and placed on an orbital shaker for 48h. The samples were then washed twice with 50 mL of 1X PBS for five minutes, before scanning. In cases where the stain did not sufficiently diffuse in the sample to visualize the fascicles, staining was repeated for an additional 48 hours. If the sample failed to stain during the additional 48h, it was stained with 1% Lugol’s Iodine, to allow for fascicular visualization; this occurred only in one sample (S8).

#### Micro-Computed Tomography Parameters

Scanning was performed using a Scanco µCT 100. The sample was glued to a 3D printed polylactic acid (PLA) board using a polyurethane adhesive (Gorilla Glue) and placed in a tube provided by Scanco. The board had 5mm spaced indentations, which allowed for coregistration of repeated scans., and histology scans were first performed with a 90 kV tube voltage, 4 µA tube current, 0.5 mm aluminum filter, 3000 projections across 360°, and reconstructed with an isotropic voxel size of 4.9 µm. A lower resolution scan initially confirmed that the stain diffused throughout the entire cross section of the nerve. In n=3 trials (‘Micro-CT rescan’ in Table 1), microwires were removed after the initial scan and the samples re-scanned at 55 kV using the same settings. This reduced voltage increased the contrast between the perineurial border and electrode tracts, but was not used with samples containing microwires because of the risk of pixel saturation with high radiopacity of the wire. All other parameters of these scans remained the same, including the resulting resolution. A small amount of phosphate buffered saline (PBS) placed at the bottom of the scan tube prevented excessive sample drying during scanning.

#### Micro-computed Tomography Analysis

*Electrode Placement:* Each nerve scan was reconstructed with a Z axis resolution of 25 um. The image intensity range was adjusted so that the dimmest voxel was 0. For placement analysis, each scan was transformed via rotation or reversal if necessary to align the superficial, implanted surface of the nerve as “up” and to make the proximal to distal directions consistent between samples, with slice 1 being the most proximal.

FIJI enabled characterization of the placement of the MiiNS contacts relative to the fascicles and their distribution throughout the nerve cross-section[35]. The 3D coordinates that defined the location of each MiiNS contact were determined by first tracing the microwire’s trajectory through the slices of the micro-CT. The slice containing the proximal appearance of the microwire, corresponding to the apex of the hook was saved for further analysis (Fig. 3). In each of these saved slices, the entire nerve cross-section was traced, and the centroid of the traced area was calculated and defined as the origin, *p*_*centroid*_ (Equation 1a). The position of the MiiNS contact was defined as *p*_*elec*_ (Equation 1b). The vector between these points was 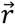,with a magnitude of *R* (Equation 2a, b). 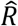 is the normalized magnitude value, determined by dividing *R* by the radius of a circle with an area equivalent to the traced nerve (Equation 3). Angle *θ* was calculated as the polar angle of vector 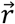 spanning from 0-360° (Equation 4).

**Figure 3.**
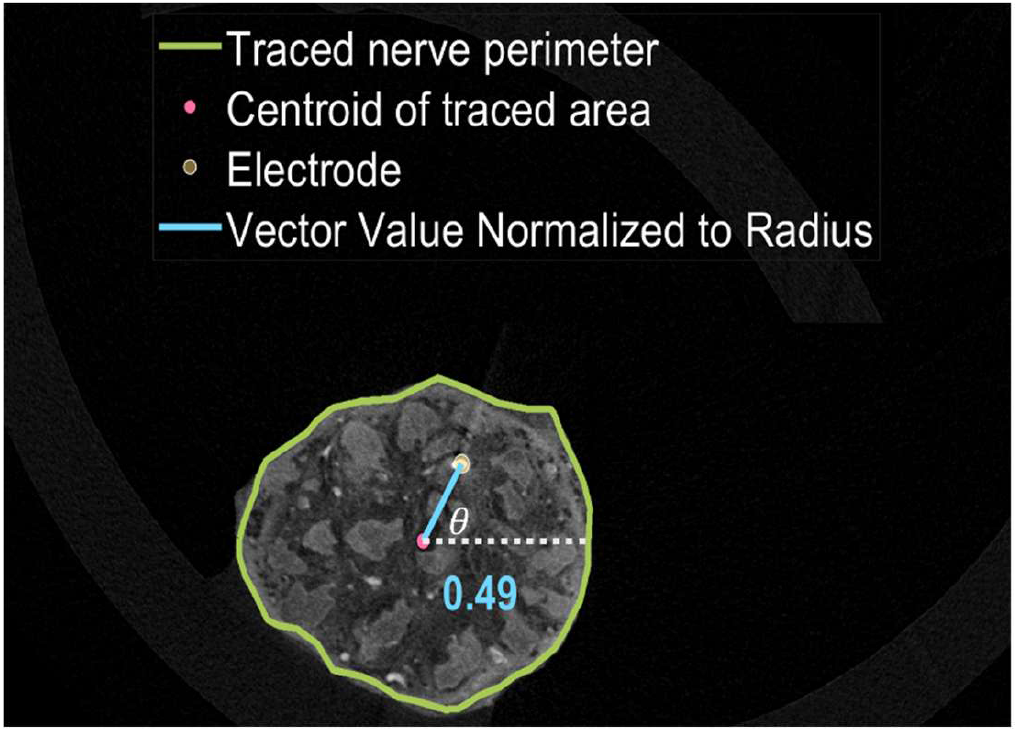
Representative micro-CT nerve slice containing the most proximal appearance of the apex of a hooked microwire contact, p_elec_ (translucent copper circle). Nerve perimeter is manually traced in green and calculated p_centroid_ is denoted in pink. Vector r (blue) with normalized magnitude 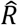 and angle θ is defined between the centroid and the electrode, considering the centroid as the origin. The normalized 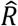 is displayed.

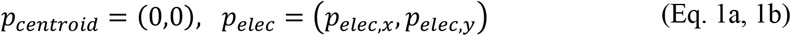

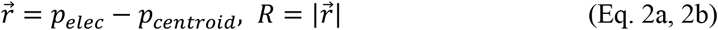

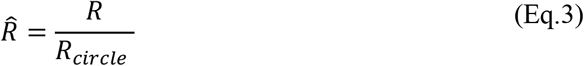

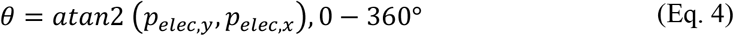

To consider the cumulative placement of all the contacts across the nerve samples in the longitudinal z direction, the midplane between the most distal and most proximal contact was centered at zero. In some cases, the normalized 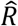 resulted in a value slightly greater than 1, which is an artifact of approximating the nerve as a perfect circle despite its oblong cross-sectional appearance in some areas. While these contacts appeared to reside outside of the nerve volume after normalization, visual inspection confirmed that these instances were within the bounds of the nerve.

#### Intraneural Placement Classification

Each microwire contact was also classified as interfascicular or intrafascicular, relative to the fascicular anatomy. Classification was based on visual inspection of each microwire’s position relative to fascicular boundaries across all visible slices and fascicles, with intrafascicular placement defined as any instance of wire traversal through or within a fascicle. The fascicles frequently demonstrated significant deflection and deformation around a microwire. Microwire placement was considered intrafascicular when the fascicle boundary was distinctly resolved, encircling the microwire. An example of this is provided later in the results. This ensured that the electrode was within the fascicular tissue rather than merely alongside it.

#### Angle of Insertion

Analysis was performed on the micro-CT scans to understand the relationship between angle of micro cannula insertion into the nerve intraoperatively and the placement of the microwire. The angle of the wire trajectory was assessed by first finding the centroid of the nerve area in all sections (FIJI, Analyze Particles)[35]. The wire position was manually tracked and segmented from its most proximal appearance to the point where it exited the nerve. Care was taken to only consider points inside the nerve, stopping segmentation upon the exit of the microwire from the nerve. The location of the apex of the microwire hook and the last point before it exited the nerve were used to define a wire trajectory vector, whereas the centroid points were linearly fit to find a nerve centroid vector, which represents the nerve’s longitudinal axis. The two-point wire fit, as opposed to fitting the entire wire, was chosen to account for the bowing that occurred in the wire, likely upon microcannula removal during implantation and would otherwise distort the wire’s true trajectory. The wire trajectory angle of each microwire was defined as the angle between the wire trajectory vector and nerve centroid vector.

#### Histology

In a subset of the nerve samples (n=4) histology was performed after micro-CT to validate the positional information content and fascicle boundaries of the micro-CT analysis (Histology’ in Table 1). In these cases, the wires had not been previously removed for rescanning. A 15 mm segment of the nerve containing the majority of the microwires was excised, and the microwires were gently removed with forceps. The position of the excised tissue was measured relative to the most proximal aspect for subsequent coregistration with prior micro-CT scans. The proximal/distal and superior/inferior orientations of the tissue were marked with tissue dye. Nerves were processed using a Pegasus Automated Tissue Processor (Leica Biosystems), which first dehydrated nerve samples with increasing concentrations of ethanol, then washed with xylene, followed by a final step of heated and vacuumed paraffin for impregnation of tissue. After processing, each nerve was divided into 3–4 tissue segments, embedded in paraffin using a Sakura TEC 6 Embedding Console, and sectioned with a Leica Histocore Autocut microtome. Every 100µm, three 4µm thick slides from each section were produced. Hematoxylin and Eosin staining was performed using an automated slide stainer (Leica Spectra ST) and Statlab Select stains. Each histological slice was coregistered with its corresponding micro-CT slice using manual inspection and matching dynamic anatomical landmarks. Micro-CT images were rotated to reflect the orientation of the histological slices. Examples of anatomical landmarks that enabled coregistration included the relative positions of dynamically merging/diverging fascicles, positions of microvasculature, positions of microwires or microwire voids and fascicular shape.

## Results

### Analysis of MiiNS Contact Placement

#### Spatial Distribution of Microwires

Each micro-CT slice containing the apex of each hooked microwire was considered for placement analysis. Each nerve sample showed spatially distinct contact positions (Fig. 4a), with average pairwise Euclidean and transverse distances across all contacts of 5352.80 ± 4049.70 µm and 1415.84 ± 753.98 µm, respectively. When all placements were considered cumulatively across the samples (Fig. 4b), the average normalized vector distance from the nerve centroid to the electrode 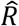 was 0.4811 ± 0.28. Contacts were placed in the central half of the nerve volume (R<0.707) 79% of the time. The mean angle (*θ*) of the vector R was 199.5± 72.2º (circular mean and StDev). To further understand the distribution of *θ*, Rayleigh’s test of uniformity was performed on the cumulative data set, which tests if the angles depart from a uniform distribution about a circle in a unimodal, von Mises distribution[36]. The cumulative dataset of placements failed to reject the null hypothesis (p=0.094), suggesting that the data did not show a statistically significant departure from a uniform distribution. The contact spacing for each sample was evaluated by calculating the Euclidian and transverse distances between each contact and all other contacts, and each contact and only its nearest neighbor (Fig. 5). On average, the Euclidian distance from any contact to its nearest neighbor was 2256.26±1760.28 µm across all samples.

**Figure 4a.**
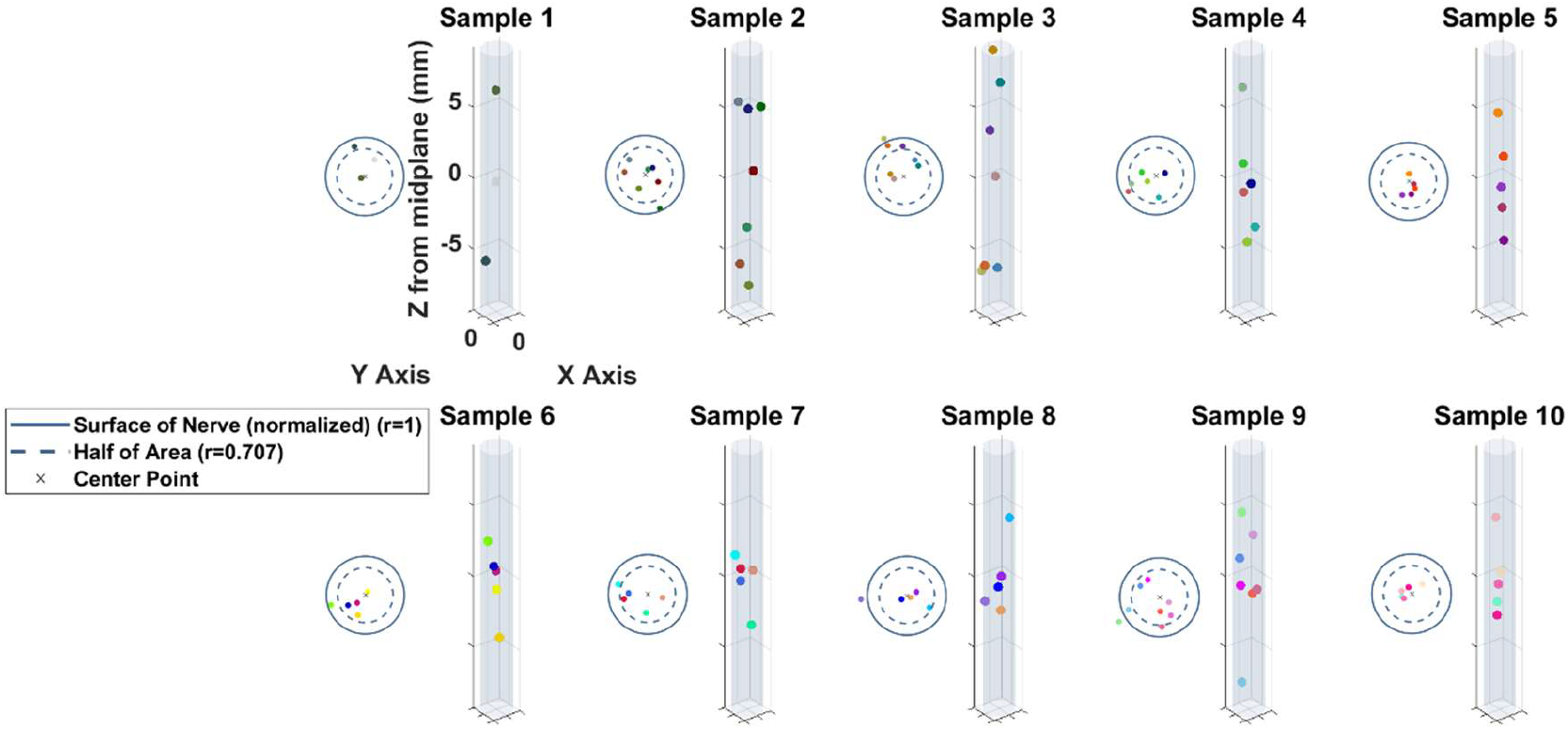
Normalized transverse (left of each nerve) and longitudinal positions of contacts in each sample (S1-10). Most contacts are placed in the center half of the nerve volume, denoted by the dashed circle on the transverse view. Each sample demonstrates a unique distribution of contacts in both the transverse and longitudinal directions.

**Figure 4b.**
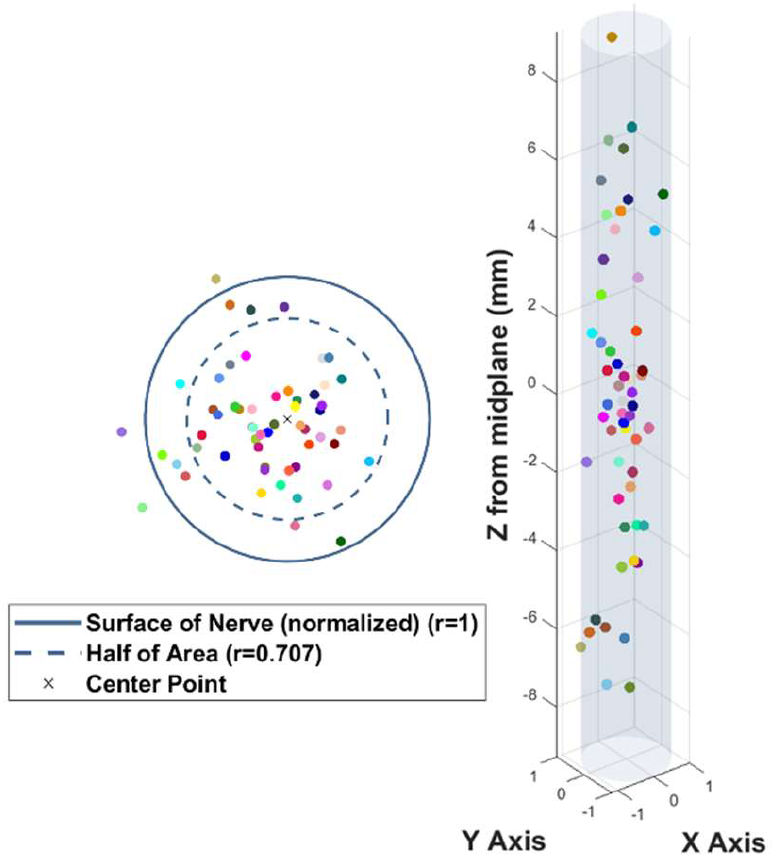
Transverse (left) and longitudinal view (right) of positions of all microwire contacts (n=56) in all nerve samples. Each contact position is a unique color, which corresponds to figure 4a. Contacts display distributed positions in terms of depth and angle in both the transverse and longitudinal directions across all the samples combined.

**Figure 5.**
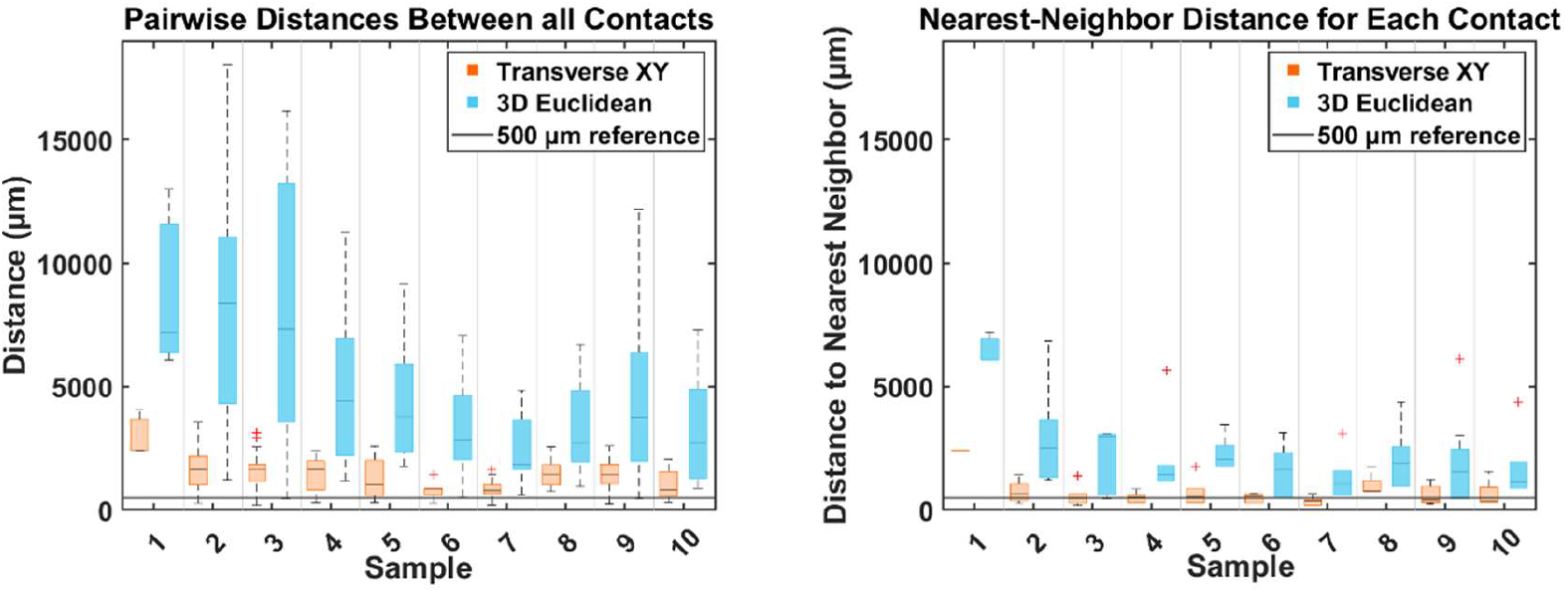
Average distances between all contact pairs (left) both in the 3D Euclidian space(blue) and in the transverse plane only (orange) across all samples (S1-10). Average distance to each contact’s nearest neighbor (right) across samples demonstrates that transverse spacing is comparable to 500um reference line. Increased contact spacing in earlier samples (S1-4), was due to repeated attempts at insertion, without the use of the tool set.

The transverse (XY-plane) nearest-neighbor distance was calculated (730.71 ± 564.83 µm) to enable comparison with other transverse electrode configurations. Also, although fascicles merge and diverge along the nerve, functionally related fibers tend to remain grouped somatotopically, making transverse spacing a meaningful indicator of potential selectivity[37].

In the longitudinal direction, the average total span between the most proximal and distal contacts within a nerve was 10.3± 4.3 mm. In context, this value is similar to the width of commonly used extraneural cuffs such as the CFINE which has a typical length along the nerve of 7 mm, or 12mm with the addition of an anode strip[6,38].In contrast to the transverse placement, the longitudinal placement was a function of the surgeon’s decision of where to place the inserter along the nerve. The surgeon aimed to place the contacts in unique locations along the nerve, within the window of exposure but was given no further specific instructions on longitudinal placement.

#### Classification of Interfascicular or Intrafascicular Placement

Each microwire’s entire trajectory within the neural tissue was classified as interfascicular or intrafascicular based on the wire’s position relative to the boundaries of fascicles (Fig. 6). In cases of intrafascicular placement, the fascicular boundary could clearly be resolved around the microwire. Of 56 total placements, n=9 showed intrafascicular behavior on any part along the wire trajectory, while the remaining 47 (84%) appeared between and alongside the fascicles. Perineurial traversal occurred in superficial and deep fascicles and was not necessarily at the site of the hooked contact. All intrafascicular placements were observed in the median nerve, and none were observed in the ulnar nerve.

**Figure 6.**
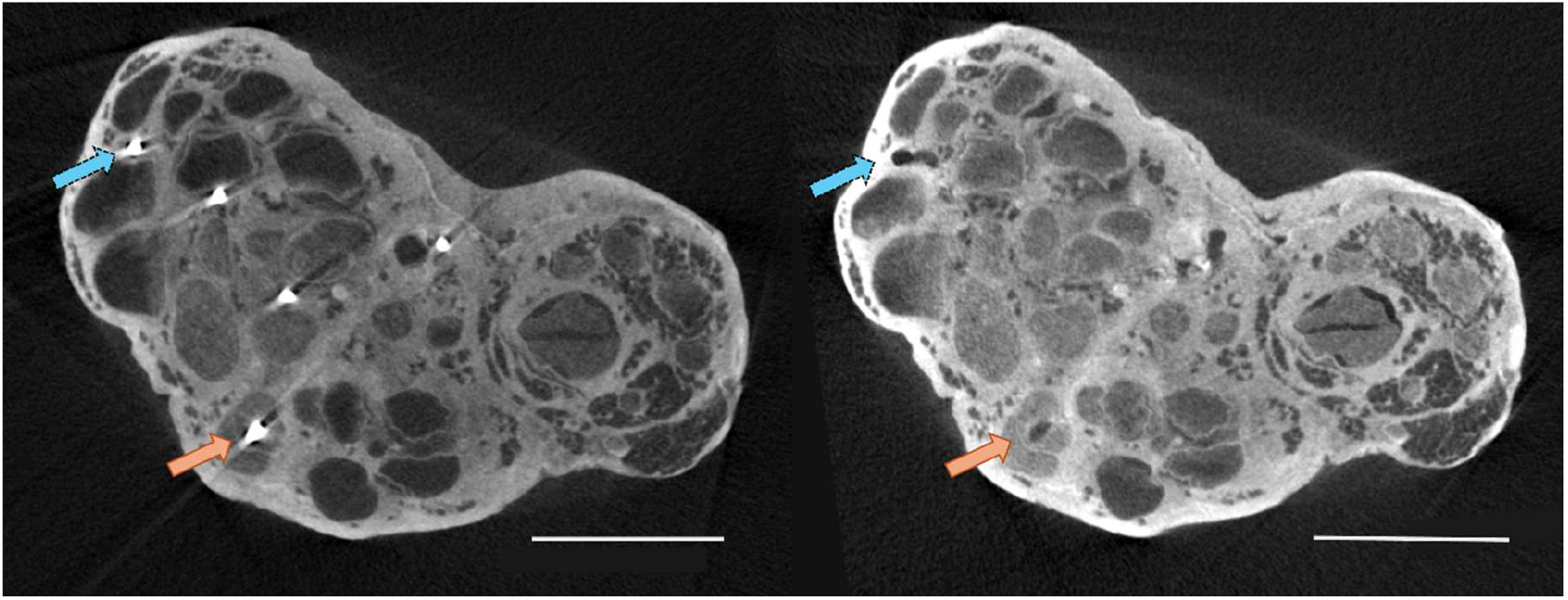
Coregistered micro-CT slices of the median nerve (S4) before wire removal (left) and after wire removal and rescanning (right). After removal, voids indicating the wires’ previous positions were apparent in most slices. Orange solid arrows indicate an example placement classified as intrafascicular, demonstrating clear traversal through a fascicle. Blue dashed arrows highlight an example interfascicular placement, barring traversal through a fascicle in any other slice along the wire’s length. Each microwire’s position was characterized along its full trajectory through the nerve. In this representative slice, only the orange arrow contact would be considered intrafascicular.

#### Role of Novel Tool Development on Electrode Placement

Without the novel tool set, the by hand insertions of S1-4 resulted in 74% interfascicular placement across 23 contact placements. The later samples (S5-10) with the novel tools were 88% interfascicular across 33 contact placements.

#### Effect of Angle of Insertion on Contact Placement

To understand how insertion trajectory of the microwire into the nerve influences the electrode’s placement within the nerve, we quantified the angle between each microwire’s trajectory and the longitudinal axis of the nerve (Fig. 7). All the wires exhibited some degree of curvature within the nerve, likely because of the retraction of the microcannula during implantation and due to the flexibility of the microwire and surrounding tissue. Quantifying the microwire’s position relative to the nerve’s longitudinal axis enabled comparison of trajectory angles both between nerve types (median vs. ulnar) and between placement types (intrafascicular vs. interfascicular). A non-parametric Wilcoxon Rank Sum Test compared the angles of wire trajectories between these groups. A Holm-Bonferroni correction adjusted for repeated comparisons[39]. Angles differed significantly across three comparisons: median versus ulnar nerves, intrafascicular versus interfascicular placements overall, and intrafascicular versus interfascicular placements within median nerves alone. Median nerves had a steeper average insertion angle, while ulnar nerve insertions were more parallel (19.0± 13.0 and 6.3± 2.3 respectively). Across all nerves, all intrafascicular insertions had a steeper angle of insertion compared to all interfascicular insertions (29.5± 20.9 and 12.9± 8.2 respectively). Finally, in the median nerve alone, where all intrafascicular placements occurred, there was a significantly steeper angle of insertion for intrafascicular placements compared to interfascicular placements (29.5± 20.9 and 16.0± 8.1 respectively).

**Figure 7.**
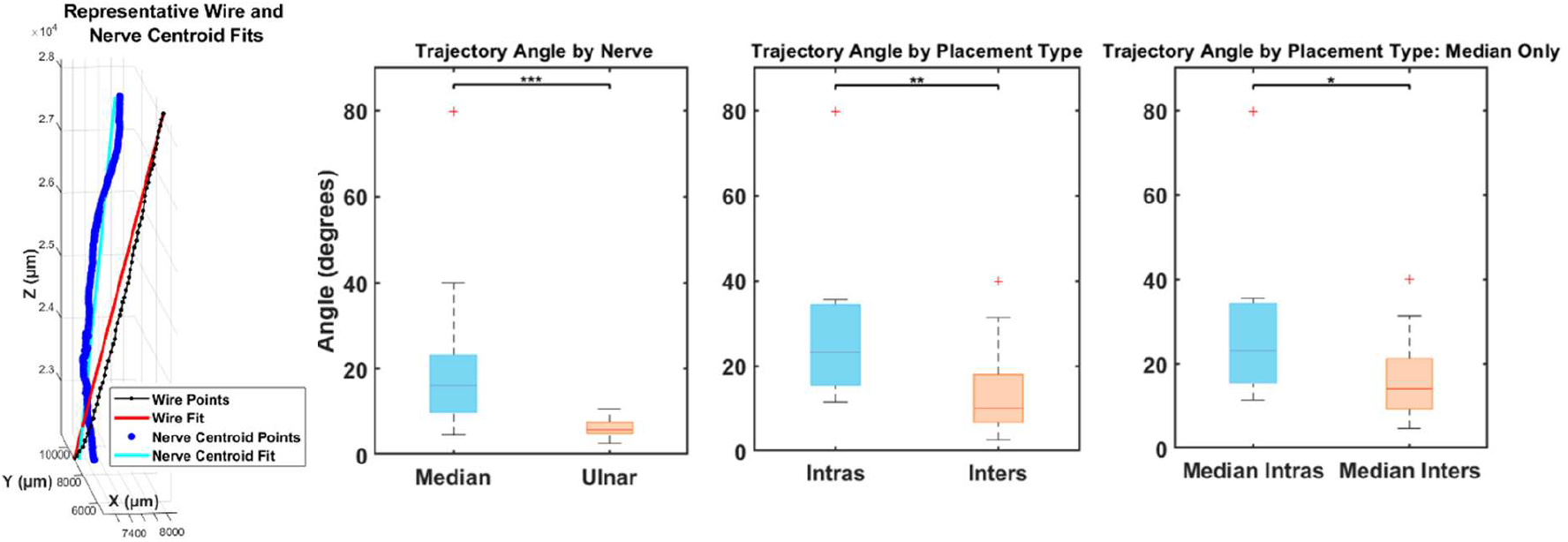
Calculation of wire trajectory angle (left). Nerve centroid (dark blue), wire trajectory (dotted black) and their respective linear fits (cyan, red). Wire trajectory angle is calculated as the angle between these two fit lines. Comparison of wire trajectory angle, and various placements is shown in the box plots. Left panel shows the comparison of the angle of wire trajectory between all insertions of median contacts (n=41) versus all insertions of ulnar contacts (n=15). Middle panel compares all instances of intrafascicular placement (n=9) to all instances of interfascicular placement (n=47), regardless of nerve type. Right most panel compares intrafascicular (n=9) and interfascicular (n=32) placements only considering placements into the median nerve. Three Wilcoxon rank-sum tests were performed, with a Holm-Bonferroni correction for family-wise error. *** indicates p<.001, ** indicates p<.01, * indicates p<.05.

#### Comparative Histological Analysis of Contact Placement

Each histological slice of the four tested nerve samples (S5,S6,S7 and S8) was successfully coregistered to its corresponding micro-CT slice using anatomical landmarks. Histological analysis was performed on samples that did not undergo repeated scanning and still contained microwires. Throughout the histological samples, the perineurial membrane was clearly visualized around each fascicle. The location of the removed wires was identified by the presence of voids and disruption of the surrounding tissue (Fig. 8). Both micro-CT and histology showed deflection of a fascicle around a microwire. When such deflection was captured by histology, the perineurial membrane was visually intact (Fig. 8, right). In all cases in which histology was available, micro-CT and histological assessments of microwire placement agreed in terms of wire position relative to the fascicles. There was no evidence of perineurial penetration in any histological sample. One of these samples exhibited intrafascicular placement on the micro-CT (Sample 5); however, the exact position of perineurial rupture was not captured histologically.

**Figure 8.**
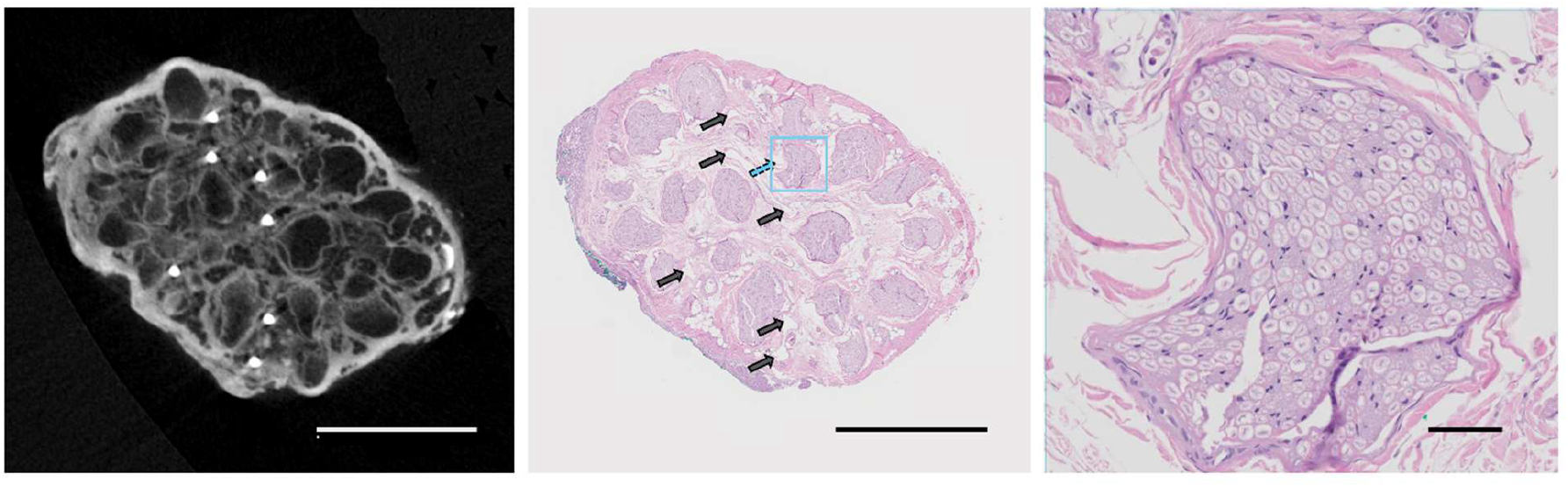
Representative coregistered micro-CT and histology slices from Sample 6. Scale bars represent 1mm (left and middle), and 50um (right). Micro-CT slice has been rotated and enlarged to match histology orientation. Black arrows are added to indicate previous location of microwires, identified as voids in the neural tissue. Some microwires deflected the shape of the fascicle significantly without penetration of the perineurium, as shown by a continuous layer of perineurial cells around the fascicle. This effect was seen several times with histology-a representative example is shown as a blue dashed arrow and is enlarged from the blue box in the right panel.

#### Surgical Tooling Assessment for Interfascicular Microwire Implantation

The combined use of the Neuroretractor, spring-loaded inserter, and gel-tipped pressure applicator enabled implantation of microwire contacts into the nerve and suggested reduced insertion time and implantation failures.

Timing data was recorded during five nerve implantations (n=25 contacts in total) with the novel toolset, and during four nerve implantations using only manual insertion (n=24 contacts in total). In the hand insertion trials, the insertion of MiiNS took 3.07± 1.2 minutes per contact, with 19/24 (79%) insertions successful on the first attempt. The most common failure mode occurred when the microwire was withdrawn from the nerve along with the microcannula, which required reinsertion (n=3). Other less common failure modes included needle buckling (n=1) and the wire falling out of the nerve before insertion (n=1). Using the novel toolset, the average insertion time per contact was 1.8 ± 0.18 minutes with 22/25 (88%) of insertions successful on the first attempt. The modes of failure were similar, with one instance of the microwire being withdrawn with the microcannula and another instance of the wire falling out of the nerve before insertion. There was an additional failure of the needle traversing through the entire nerve (n=1).

## Discussion

Micro-CT analysis showed contacts that were well distributed throughout the nerves cross section and length. The MiiNS contacts were distributed throughout the central half of the nerve. This provides access to areas of the nerve that are difficult to selectively activate with circumneural electrodes, while still not penetrating the perineurium. The polar angles of the electrode contacts were not confined to a single plane within the nerve. Instead, they spanned the full circumference, and statistical analysis showed no significant deviation from a uniform polar distribution. This test indicates that the electrode contacts were well distributed radially. This configuration is different from existing flexible intrafascicular electrodes that implant contacts in a single plane, and should offer expanded stimulation access[12,40]. Previous modeling studies of both extraneural and intrafascicular interfaces suggest that transverse spacing between electrode contacts should be approximately 0.5–1 mm to optimize selectivity[41,42]. Existing devices fall within this range, including the circumneural CFINE which has 500µm contact spacing, and intrafascicular configurations such as the TIME which uses 750µm contact spacing[38,41]. The MiiNS transverse contact spacing also falls within this optimal range, supporting the ability to target distinct axonal populations effectively. Furthermore, the longitudinal placement of MiiNS contacts is easily customizable. Contacts can be spread along a nerve or clustered in a very limited insertion window, depending on the application, or the nerve anatomy. In future use, MiiNS could function as a standalone interface or as an augmentation strategy to other interfaces to access deep or distributed axonal populations, which is a remaining research question.

Micro-CT analysis also showed that most of the microwire contacts were placed between the fascicles and did not traverse into a fascicle at any point along its trajectory. This is an expected result due to the use of a blunt microcannula and the biomechanical properties of the perineurium. By placing electrodes between the fascicles, we hypothesize that we limit violations to the blood nerve barrier, and in turn reduce neural damage. However, future chronic studies are needed to quantify the chronic nerve response to MiiNS and investigate this claim. Histological evidence corroborated the relative positions of the microwires, and fascicles shown on micro-CT. Deflection of the fascicular boundary was commonly observed in micro-CT data. In such cases, metal artifacts and the absence of distinct cellular contrast made it difficult to delineate perineurial boundaries. Spatially registered histological sections in several of these cases revealed an intact perineurial membrane. This finding indicates that the fascicle is capable of conforming to either the insertion cannula or the implanted microwire. Importantly, deformation around the electrode does not necessarily indicate membrane rupture, and the perineurium can remain structurally intact despite this local displacement. One caveat to this claim is that systemic perfusion of fixative was not performed prior to euthanasia, and we cannot be sure that the deformation is not simply an artifact of the loss of fascicular pressure postmortem.

The angle of wire trajectory of the MiiNS microwires demonstrated a significant effect on intrafascicular or interfascicular placement in the nerve as shown by post hoc analysis. Specifically, steeper wire trajectory angles were associated with a higher incidence of intrafascicular electrode placement. Unexpectedly, all intrafascicular placements occurred in the median nerve. One possible explanation for this difference was that the median nerve was observed to have a different surgical exposure, resulting in the need to hold the inserter at a steeper angle during implantation. Although anatomical differences may also have contributed to the higher incidence of intrafascicular placement in the median nerve, analysis of median nerve samples alone demonstrated that intrafascicular wires followed significantly steeper trajectories than interfascicular insertions. This finding supports that the effect was related to insertion angles rather than just nerve-specific anatomy, but further study is needed to elucidate this relationship. The role of insertion angle of peripheral nerve interfaces has practical implications for those who are intentionally trying to target intrafascicular placement, or for those aiming to avoid perineurial penetration. Future designs for nerve implants should consider controlling the angle of insertion to target electrode placement relative to the neural tissues.

Specialized tools are important for quick and controlled placement of interfascicular electrodes. To minimize neural damage from mechanical factors, interfascicular nerve interfaces must be small and flexible. The insertion of these flexible interfaces into the nerve necessitates a rigid insertion guide or cannula. However, nerve mobility and space limitations render insertion with an insertion cannula alone untenable. We engineered a system of tools that enabled rapid placement of microwire electrodes within the nerve. The Neuroretractor, spring loaded inserter and gel tipped applicator facilitate successful placement of 50 µm microwires which would readily buckle upon contact with soft tissue. These tools are agnostic to the material and configuration of the contacts. Microwire contacts are delivered in a hooked configuration, and do not require passing through the entire nerve, and do not require sutures. In addition, the distally hooked delivery configuration allows for the upstream lead body and connectors to have any configuration, as they do not pass through the micro-cannula during implantation. In this study, the microwire interface largely served as a proof of concept for the placement of interfascicular electrodes. However, future iterations may include multi-contact insertions, smaller or more flexible interfaces, or mechanically dynamic interfaces. Implantation of MiiNS was successfully performed in a large-animal surgical setting, demonstrating the feasibility of this approach in nerves that are anatomically and clinically relevant to humans. In addition, these tools – particularly the spring-loaded inserter – are amenable to development of a percutaneous approach in the future. A percutaneous approach could further reduce the surgical burden, while maintaining selectivity for a range of clinical applications.

Interfascicular interfaces are an emerging class of interfaces for neuroprosthetic peripheral nerve stimulation. Compared to other interface designs, these electrodes occupy middle ground. Like intrafascicular configurations, interfascicular electrodes are placed within the nerve, allowing distributed access to target axonal populations, especially at the center of the nerve. Unlike intrafascicular electrodes, interfascicular electrodes aim to minimize traversal of the protective perineurium. In turn, we hypothesize this will reduce detriment to chronic nerve health. While circumneural electrodes offer stable and selective stimulation, the central population of axons remains challenging to selectively stimulate in large nerves. Thus, interfascicular electrodes have potential to combine benefits of increased access to axon populations with fewer instances of disruption to the perineurium.

### Limitations

Micro-CT with phosphotungstic acid (PTA) staining enables high-resolution, three-dimensional, continuous visualization of peripheral nerve anatomy, as shown here and in the literature[43]. However, this technique is limited in its ability to definitively resolve the boundaries of individual fascicles at very high resolution, especially in the presence of metal artifacts. As such, classifications of interfascicular versus intrafascicular placement based solely on micro-CT data should be interpreted with a degree of caution. Histology complements micro-CT analysis by confirming tissue identity and demonstrating intact perineurium, but it requires removal of the electrode and discrete sectioning of the tissue. The histology presented here was able to identify electrode tracts at discrete locations along the nerve but could not conclusively determine whether the microwires violated the perineurium along their entire length. Further, a positive case of perineurial membrane rupture was not sampled with histology, despite sampling a nerve that exhibited potential intrafascicular placement in micro-CT. This means that positive markers of penetration of the perineurium on micro-CT could not be validated against histology. Despite this, without micro-CT analysis, it would have been difficult to identify traversals as obtaining a specific cross-section at the point of traversal is not guaranteed.

The angle of insertion into the nerve was not controlled for in the experimental design and was instead examined post hoc. Therefore, a further examination of the relationship between the angle of insertion and penetration through the perineurium would be beneficial. Understanding how the angle of insertion may impact the force required to penetrate the epineurium and perineurium would be helpful to guide peripheral nerve interface designs and was not explored here.

Future chronic studies will be important to resolve these uncertainties and to understand the overall feasibility of MiiNS. Chronic implantation will also cause recruitment of fibrotic encapsulation tissue around microwires, which will leave distinct tracts for better histological and possible micro-CT placement identification in the future. In addition, the chronic electrical stability and stimulation capability of these electrodes remains a future avenue of study.

## Conclusion

Micro-CT characterization of acute electrode placement in a large-animal porcine model demonstrated the feasibility of interfascicular electrodes as a means of balancing the trade-off between interface selectivity and invasiveness. MiiNS demonstrated the distributed placement of microwires into the interfascicular space, while minimizing traversal into the fascicles. The complete trajectory of each microwire contact revealed that angle of insertion may play a role in achieving interfascicular placement. This insight will inform future designs and could be useful for interfascicular and intrafascicular implantation strategies alike. Chronic study of the long-term impacts of MiiNS on nerve health are needed before to translation. However, we believe that MiiNS may be a promising strategy to reduce interface invasiveness while maintaining high spatial selectivity by accessing hard-to-reach central axonal populations in the nerve. The combined potential for percutaneous implantation and increased access to the nerve could yield increased therapeutic capacity of neuromodulation across nerve targets and patient populations.

## Acknowledgments

We would like to thank the veterinary staff at Case Western Reserve University’s Animal Resource Center for assisting in animal experiments and the management of regulatory compliance. We also thank J. Dunning for support in facilitating engineering efforts, A. Upadhye for running micro-CT samples, J. Coleman for histology support and E. Imka for graphic design.

## Data Availability Statement

The data cannot be made publicly available upon publication because they are not available in a format that is sufficiently accessible or reusable by other researchers. The data that support the findings of this study are available upon reasonable request from the authors.

## Ethics statement

The surgical and experimental procedures were performed with the approval and oversight of the Case Western Reserve University, Institutional Animal Care and Use Committee to ensure compliance with all federal, state and local animal welfare laws and regulations.

## Author Contribution

M.A.R. led data collection, data analysis and manuscript composition. V.I.M. contributed to data collection. C.T. developed the micro-CT protocol with M.A.R. and contributed to manuscript composition. A.J.S. advised on methodology and manuscript composition. J.Z.B. gave input on tooling design, facilitated the surgical aspects of data collection and advised on manuscript composition. D.J.T. led ideation for the study, and advised on experimental design, data analysis and manuscript composition.

